# Characterizing the Dynamical Complexity underlying Meditation

**DOI:** 10.1101/521963

**Authors:** Anira Escrichs, Ana Sanjuan, Selen Atasoy, Ane López-González, César Garrido, Estela Càmara, Gustavo Deco

**Affiliations:** Theoretical and Computational Neuroscience Group, Center for Brain and Cognition, Department of Information and Communication Technologies, Universitat Pompeu Fabra, Barcelona, Spain; Cognition and Brain Plasticity Unit, Bellvitge Biomedical Research Institute (IDIBELL), L’Hospitalet de Llobregat, Barcelona, Spain; Department of Psychiatry, University of Oxford, Oxford, England; Radiology Unit, Hospital Clínic Barcelona, Barcelona, Spain; Department of Cognition, Development and Educational Psychology, University of Barcelona, Barcelona, Spain; Institució Catalana de la Recerca i Estudis Avançats (ICREA), Barcelona, Spain

**Author notes:** Corresponding author; Universitat Pompeu Fabra, C Ramon Trias Fargas, 25-27, Barcelona, 08005, Spain,; Universitat Pompeu Fabra, C Ramon Trias Fargas, 25-27, Barcelona, 08005, Spain.

**Keywords:** ignition, whole-brain, meditation, resting-state, fMRI, integration, dynamical complexity, brain-states

## Abstract

Over the past 2,500 years, contemplative traditions have explored the nature of the mind using meditation. More recently, neuroimaging research on meditation has revealed differences in brain function and structure in meditators. Nevertheless, the underlying neural mechanisms are still unclear. In order to understand how meditation shapes global activity through the brain, we investigated the spatiotemporal dynamics across the whole-brain functional network using the Intrinsic Ignition Framework. Recent neuroimaging studies have demonstrated that different states of consciousness differ in their underlying dynamical complexity, i.e., how the broadness of communication is elicited and distributed through the brain over time and space. In this work, controls and experienced meditators were scanned using functional magnetic resonance imaging (fMRI) during resting-state and meditation (focused attention on breathing). Our results evidenced that the dynamical complexity underlying meditation shows less complexity than during resting-state in the meditator group but not in the control group. Furthermore, we report that during resting-state, the brain activity of experienced meditators showed higher metastability (i.e., a wider dynamical regime over time) than the one observed in the control group. Overall, these results indicate that the meditation state operates in a different dynamical regime than the resting-state.

## Introduction

During the last 2,500 years, contemplative traditions have explored the nature of the mind through self-discipline and self-observation. Meditation per se is not a philosophy or a religious practice, but a method of mental training which enables to cultivate a variety of human abilities, ranging from developing a clearer mind, enhancing attention to cultivating altruistic love and compassion towards other beings [1].

In the last decade, fMRI studies exploring the neural correlates of meditation have revealed important insights into how this mental training changes brain function and structure [2, 3, 4, 5, 6, 7, 8, 9, 10, 11, 12, 13]. Yet, little is known about how meditation influences the capability to transmit information across the whole-brain functional network.

Recently, it has been proposed that a brain state could be defined by measuring how the broadness of communication is elicited and distributed through the brain over time, i.e by characterizing its underlying dynamical complexity [14]. Investigating the propagation of the neural activity by measuring their dynamical implications [15] across the whole-brain network may help to explain the fundamental principles of the underlying mechanisms of different brain states [16, 17, 18, 19]. Theoretical methods have been successfully applied to characterize different states of consciousness such as wakefulness, sleep, anesthesia or psychedelic states [20, 21, 22, 23, 24, 14].

Here, we investigate the brain’s macro-scale mechanisms underlying meditation as well as meditation-induced long-term changes in resting-state using the Intrinsic Ignition Framework [25, 14]. This data-driven method allows studying the spatiotemporal dynamics across the whole-brain functional network by measuring the effect of naturally occurring local activation events on whole-brain integration.

## Methods

### Participants

A total of forty participants were recruited for this experiment. Half of the participants were experienced meditators (mean (SD) age = 39.8 (10.29); education years = 13.6; mean (SD) hours meditation experience = 9526.9 (8619.8); 7 females) and were recruited from Vipassana communities of Barcelona. All of them had a minimum of 1,000 hours of meditation experience and confirmed that they maintained daily practice (>1 hour/day). The other half were well-matched control participants with no prior meditation experience (mean (SD) age = 39.75 (10.13); education years=13.8; 7 females). No significant differences in terms of age, educational level and gender were found between groups. Participants reported no history of neurological disorder, provided written informed consent, and were compensated for their participation. The study was approved by the Ethics Committee of the Bellvitge Hospital in accordance with the Helsinki Declaration on ethical research.

### Resting-state and meditation fMRI

A total of 450 brain volumes in each condition were analyzed (≈ 15 min). During rest, participants were asked to look at a cross fixated on the screen, not thinking anything in particular, remain as motionless as possible and not falling asleep. After resting acquisition, all participants were engaged in meditation. Meditators were asked to practice anapanasati meditation (focused attention on breathing). In this type of meditation, subjects try to concentrate all their attention on natural breathing, and when they realize that the mind wanders, they must recognize it and come back to natural breathing without judgment. Controls were instructed in meditation before they had been scanned following the instructions as taught by S.N. Goenka. Controls confirmed that they understood the procedure after having done a simulation.

### MRI Data Acquisition

MRI images were acquired on a 3T TIM TRIO scanner (Siemens, Erlangen, Germany) using 32-channel receiver coil. The high-resolution T1-weighted images were acquired with 208 slices in the sagittal plane, repetition time (TR) = 1970ms, echo time (TE) = 2.34ms, TI = 1050ms, flip angle = 9°, field of view (FOV) = 256mm, voxel size 1×1×1mm. Resting-state and meditation fMRI were performed by a single shot gradient-echo EPI sequence (TR = 2000ms; TE = 29ms; FOV = 240mm; in-plane resolution 3mm; 32 transversal slices with thickness = 4mm; flip angle = 80°).

### Preprocessing

Preprocessing was computed using the Data Processing Assistant for Resting-State fMRI (DPARSF) [26]. Preprocessing included: manually reorienting T1 and EPI images; discarding the first 10 volumes due to magnetic field inhomogeneities; slice-timing correction; realignment for head motion correction; T1 co-registration to functional image; European regularization segmentation; removal of spurious variance through linear regression: six parameters from the head motion correction, the global mean signal, the white matter signal, and the cerebrospinal fluid signal, CompCor; removal of the linear trend in the time-series; spatial normalization to the Montreal Neurological Institute (MNI); spatial smoothing with 6mm FWHM Gaussian Kernel; and band-pass temporal filtering (0.01-0.25Hz) [27, 28]. Finally, we extracted the time-series according to a resting-state atlas of 268 nodes which ensures the functional homogeneity within each node [29].

One meditator was removed due to incidental findings in the MRI session, and 3 controls during meditation and 1 control during rest were excluded due to a head rotation greater than 2mm or than 2°. Moreover, the frame-wise displacement (FD) [30] was calculated due to its consideration of voxel-wise differences in its derivation [31]. Subjects with head motion greater than 2 standard deviations above the group average and movement in more than 25% of time points were excluded from analysis. FD correction led to exclusion of 1 control during meditation. Therefore, a total of 4 controls during meditation were excluded and 1 control during rest.

### Intrinsic Ignition Framework

The Intrinsic Ignition Framework [25] measures the degree of elicited whole-brain integration of spontaneously occurring events across time. Figure 1 describes the algorithm to obtain the intrinsic integration across events of each brain area. Driving events are captured by applying the method of Tagliazucchi and colleagues which measures dynamical neural events for each brain area [32]. Events are fixed as a binary signal by transforming the time-series into z-scores, *z_i_*(t), and imposing a threshold *θ* such that the binary sequence *σ*(t)=1 if *z_i_*(t) > *θ*, and is crossing the threshold from below, and *σ*(t)=0 otherwise. If a brain area has triggered an event, then the neural activity of all brain areas is measured in the set time window of 4TR. A binary matrix is obtained representing the connectivity between brain areas exhibiting simultaneous activity. Afterward, the measure of global integration [19] is applied, returning the broadness of communication across the network for each driving event (i.e the largest subcomponent). Finally, the process is repeated for each spontaneous neural event, producing the Intrinsic-Driven Mean Integration (IDMI) and the variability as the standard deviation of the Intrinsic-Driven Integration for each brain area in the whole-brain network.

**Figure 1:**
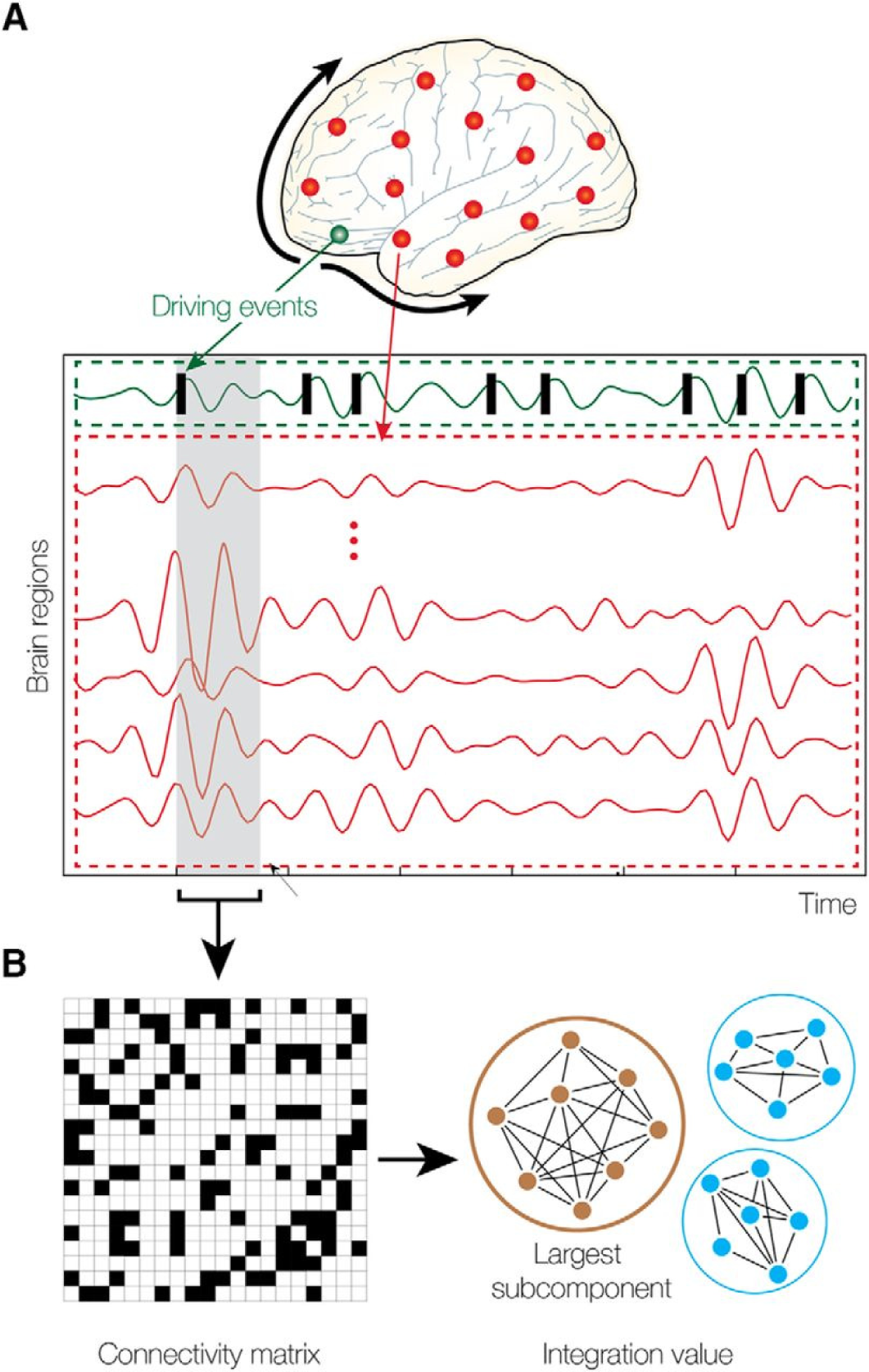
Measuring intrinsic ignition. **(A)** Events were captured applying a threshold method [32] (see green area). For each event elicited (gray area), the activity in the rest of the network was measured in the time-window of 4TR (see red area). **(B)** A binarized matrix was obtained which represents the connectivity between brain areas where activity was simultaneous. **(C)** Applying the global integration measure [19], we obtained the largest subcomponent. Repeating the process for each driving event, we calculated the mean and the variability of the Intrinsic-Driven Integration for each brain area across the whole-brain network.

### Comparisons

Statistical analyses were carried out in Matlab (R2017a). We used Monte Carlo permutation test (number of iterations 10,000) to test the differences between conditions. Furthermore, to ensure that the observed results were not obtained by chance, we applied a surrogate data testing method. We created the surrogate data using a random permutation at each time point of the original time-series and measured the ignition in each spontaneous event on the randomized data. We repeated the process 50 times and then, we compared the average randomized data to the original time-series.

## Results

### Intrinsic-Driven Mean Integration (IDMI)

We first calculated the IDMI in both states (resting-state and meditation) for each group (controls and meditators). Figure 2A shows the IDMI for each group and brain state, while Figure 2C shows the IDMI for each group and each brain area. The IDMI captures the spatial diversity as differences in average intrinsic ignition profiles across the different nodes. The brain activity of meditators during resting-state showed the highest values of the IDMI compared to the control group (p<0.001, Monte-Carlo simulations after Bonferroni correction). Furthermore, this value decreased significantly when meditators were engaged in meditation (p<0.001, Monte-Carlo simulations after Bonferroni correction). In contrast, controls did not show any differences between resting-state and meditation conditions.

**Figure 2:**
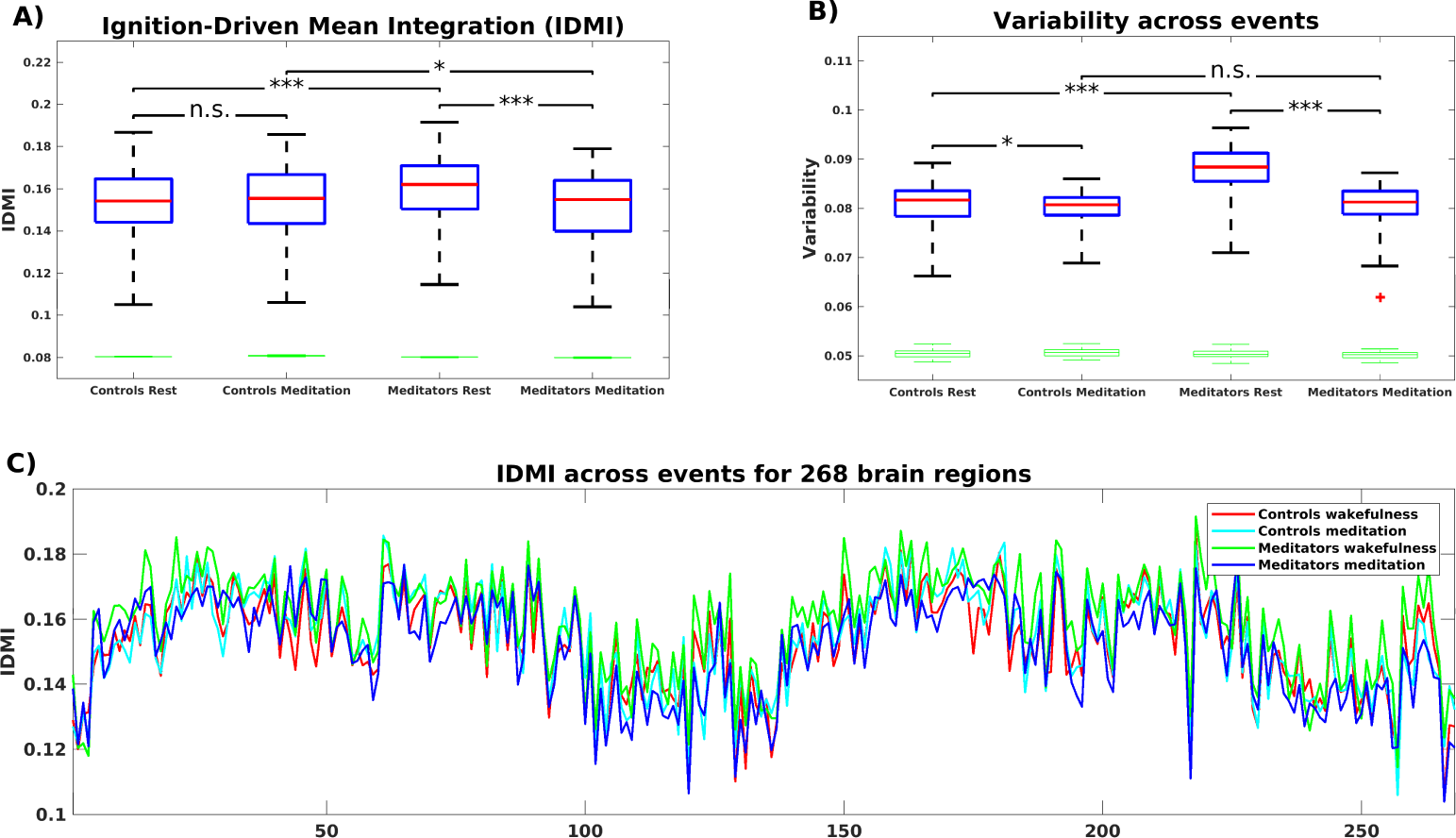
**(A)** Mean of the Intrinsic-Driven Integration (IDMI) for each group during resting-state and meditation state. The IDMI was higher in meditators than in controls during resting-state and lower in meditators during meditation. No significant differences were observed in controls between conditions. Furthermore, we show the box plot from the surrogate IDMI data (on the bottom in green). The randomized data were significantly smaller than the original time-series, showing the robust statistical comparisons. **(B)** Both controls and meditators showed higher local metastability across the whole-brain during resting-state compared to meditation. However, the effect was significantly larger for meditators. Furthermore, the metastability in resting-state was significantly higher for experienced meditators than for controls. P-values are based on Monte-Carlo simulation after Bonferroni correction, * represents p<=0.05, *** represents p<=0.001 and n.s represents not significant. **(C)** IDMI across events for each group during resting-state and meditation for 268 brain regions.

### Variability of Intrinsic-Driven Integration

Next, we calculated the variability of the Intrinsic-Driven Integration in both states (resting-state and meditation) for each group (controls and meditators). Figure 2B shows the variability for each group and brain state. The variability describes the heterogeneity of each brain area, which is closely connected to its local metastability [25]. Thus, it describes how the local activity in each brain area changes across time. High levels of metastability in a node represent a more dynamic function over time, while lower levels represent greater stability. The brain activity of controls and experienced meditators showed higher functional variability (ie, metastability) in resting-state than in meditation. Nevertheless, the effect was significantly larger for meditators (p<0.001, Monte-Carlo simulations after Bonferroni correction) than for controls (p<0.05, Monte-Carlo simulations after the Bonferroni correction). Furthermore, the metastability in resting-state was significantly larger for experienced meditators than for controls (p<0.001, Monte-Carlo simulations after the Bonferroni correction).

## Discussion

A growing scientific interest lies in the characterization of the meditation state. Hasenkamp and colleagues [5] captured the interactions between four cognitive phases during meditation but disregarding the dynamical properties, which contain relevant spatiotemporal information. Mooneyham and colleagues applied a dynamical functional connectivity approach dissociating mental states during a meditation scan. The authors reported that after a 6 weeks intervention mindfulness program, subjects spent more time in the state of focused attention and less in the state of mind-wandering [11]. In addition, a study that applied graph theoretical analysis [33] characterized the degree of the hierarchical organization during meditation. This study revealed that some nodes had the highest integration degree during rest but the lowest during meditation, and vice versa. Our work extends these findings by exploring the brain activity during meditation by characterizing the dynamical complexity in terms of how local information is broadcasted across the whole-brain.

Here, we have characterized the dynamical complexity underlying resting-state and meditation in healthy controls and experienced meditators as evidenced by the level of intrinsic ignition. Specifically, in meditators but not in controls, we observed a significant increase of intrinsic ignition during resting-state compared to meditation (Figure 2A). In addition, during resting-state, meditators showed the maximal variability of intrinsic ignition (i.e metastability) across the whole network, revealing a state of maximum network switching (Figure 2B).

Our results showing an increase of intrinsic ignition during rest compare to meditation are consistent with recent studies on information propagation across the brain. Irrmischer and colleagues found a shift from more complex brain dynamics during rest to a state of reduced information propagation during meditation, importantly, only in meditators [34]. Furthermore, Gard and colleagues demonstrated using graph theory that yoga and meditation practitioners showed greater network integration than controls during rest [35]. In addition, the increase of metastability in meditators during resting-state is congruent with the increase of the temporal complexity of oscillations during rest in meditators observed in the previously mentioned study [34]. Moreover, studies applying a dynamical functional connectivity approach found that individuals with high trait mindfulness, transitioned more frequently between brain states at rest [13, 36].

To sum up, our investigations suggest that the dynamics underlying meditation differs from the dynamics underlying resting-state. Even more, the degree of expertise in meditation of the subjects significantly changes the dynamical complexity underlying both resting-state and meditation. Future longitudinal studies (before and after an intervention in the same participants) will be needed to investigate the necessary time-training time necessary to alter the dynamical complexity induced by meditation.

## Author Contributions

AE and GD designed the study. EC and CG designed the MRI protocol. AE collected the data. AE and AS pre-processed the fMRI data. AE, AS and AL performed the analyses. AE, AS, GD and SA interpreted the results. AE wrote the first version of the manuscript. All authors contributed to manuscript revision, read and approved the submitted version.

## Acknowledgments

A.E. is supported by a Francisco J. Varela Award from the Mind and Life Europe. A.S. is supported by the Spanish Ministry of Economy and Competitiveness Grant FPDI2013-17045. G.D. is supported by the Spanish Ministry Research Project PSI2016-75688-P (AEI/FEDER), by the European Union’s Horizon 2020 FET Flagship Human Brain Project 785907 HBP SGA2, by the Catalan Research Group Support 2017 SGR 1545 and by the Foundation Marato TV3 2016.

## Conflict of Interest Statement

The authors declare that the research was conducted in the absence of any commercial or financial relationships that could be construed as a potential conflict of interest.

